# Female oviposition site selection influences hatching success and embryonic development in a Neotropical glass frog

**DOI:** 10.64898/2026.06.14.732161

**Authors:** J. Sebastián Curaca-Fierro, Johana Goyes Vallejos

## Abstract

For oviparous animals, the decision of where to lay eggs is critical, as offspring remain sessile from oviposition through hatching and are thus unable to escape unfavorable conditions. Consequently, females are expected to select oviposition sites that benefit embryo development and survival. This may be particularly relevant for arboreal frogs, which typically lay eggs on leaves overhanging water, where embryos are exposed to predation, desiccation, and other risks until hatching. Yet studies directly linking maternal substrate choice to embryo survival remain scarce. Here, we examine how oviposition substrate influences embryo survival in the Emerald glass frog (*Espadarana prosoblepon*), a species in which females deposit eggs on multiple substrates, providing a rare opportunity to test how oviposition decisions affect reproductive success. Monitoring clutches *in situ*, we compared microclimatic conditions, hatching success, and sources of embryo mortality between the most used substrates: the spike moss *Selaginella diffusa* and leaves. Additionally, we conducted a two-choice experiment in semi-captivity to test whether females preferentially select one substrate over the other. Although microclimatic conditions did not differ between substrates, hatching success was significantly higher on *S. diffusa*, which also experienced less predation. In the two-choice experiment, all females laid their eggs on *S. diffusa*, and those clutches had higher hatching success and faster embryonic development rates than those on leaves. Together, these results support the hypothesis that non-random oviposition site selection in *E. prosoblepon* is driven by the maximization of embryo survival, demonstrating that substrate choice has measurable fitness consequences for the offspring.

## Introduction

Oviposition site choice is a critical reproductive decision with direct consequences for both parental fitness and offspring survival (Buxton and Sperry 2017). In oviparous species, such as birds, reptiles, fish, amphibians, and invertebrates, females often face a trade-off between minimizing risks to themselves and their offspring and selecting sites with environmental conditions that favor successful hatching. Hence, natural selection is expected to favor non-random oviposition behavior, whereby parents select sites that provide conditions suitable for offspring survival and development (Refsnider and Janzen 2010). The quality of oviposition sites is shaped by a combination of abiotic factors (i.e., temperature, humidity, and rainfall) and biotic factors, including the presence of predators and parasites (Resetarits 1996). For example, adult Kentish plovers (*Charadrius alexandrinus*) prefer open nesting sites, which, despite greater heat stress, allow them to more easily detect and chase away egg predators (Amat and Masero 2004). Similarly, brown anole (*Anolis sagrei*) females preferentially bury their eggs in cooler, more humid microhabitats, particularly early in the nesting season, as soil humidity strongly influences embryonic development (Pruett et al. 2020). Thus, while some species can actively mitigate risk at the egg laying site through parental defense, the decision of where to lay eggs is particularly high stakes in species that provide no further care after oviposition, as offspring survival depends entirely on the conditions at the selected site.

Given the fitness consequences of selecting a low-quality oviposition site, females of many species have evolved the ability to assess oviposition site quality based on the very conditions that influence offspring survival, a behavior consistent with the ‘mother knows best’ hypothesis, which predicts that females preferentially select oviposition sites that maximize offspring survival (Refsnider and Janzen 2010). Support for this hypothesis spans diverse taxa. In herbivorous insects, eggs laid on preferred host plant species show higher survival rates (Gripenberg et al. 2010; Jones 2022). In reptiles, oviposition site selection has direct fitness consequences, as seen in female Eastern fence lizards (*Sceloporus undulatus*), where selecting nest sites with intermediate temperatures and high humidity increases hatching success (Warner and Andrews 2002). Precipitation represents another important axis of oviposition site quality. For example, in Diamondback moths (*Plutella xylostella*), females predominantly oviposit on the surface of leaves positioned lower on the host plant and along leaf veins. Experimentally simulated rainfall demonstrated that eggs in these regions are substantially less likely to be dislodged (Rahman et al. 2019). Furthermore, predation pressure constitutes perhaps the most direct selective force on oviposition site choice, given the potential for total clutch loss. Yellow warblers (*Setophaga petechia*), for example, preferentially nest in concealed microhabitats in response to high predation rates from avian predators (Latif et al. 2012). Collectively, these examples illustrate that abiotic and biotic factors are key determinants of oviposition site quality, yet how preferences for specific site characteristics translate into measurable fitness benefits remains an open question in many systems.

Arboreal frogs are particularly well-suited for addressing the question of how oviposition site characteristics influence embryo survival because their embryos develop externally on vegetation, where microhabitat conditions vary substantially and can be directly linked to offspring survival. Research in these systems has examined how abiotic and biotic conditions influence embryo survival and oviposition site choice (Buxton and Sperry 2017; Sánchez-Ochoa et al. 2020). For instance, female hourglass treefrogs (*Dendropsophus ebraccatus*) select oviposition sites that are less likely to experience dehydration or high predation risk (Touchon and Warkentin 2010; Touchon and Worley 2015). In contrast, one potentially important yet understudied dimension of oviposition site quality is the physical substrate on which eggs are deposited. Recent evidence suggests this factor can significantly influence embryo fitness: in the forest green treefrog (*Zhangixalus arboreus*), eggs laid on soil rather than on leaves experience reduced nighttime cooling and achieve higher hatching success (Ichioka and Kajimura 2024). However, testing whether females actively choose among substrates and whether such choices affect hatching success is difficult in most arboreal frogs because oviposition typically occurs on a single substrate, usually leaves. This pattern is widespread across arboreal lineages, including Hylidae (Haddad and Sawaya 2000), Hyperoliidae (Vonesh 2000), Rhacophoridae (Liao and Lu 2010; Poo and Bickford 2013), and most Centrolenidae (Delia et al. 2010; Cabanzo-Olarte et al. 2013; Williams 2024). An exception to this pattern is the Emerald glass frog *Espadarana prosoblepon* (Family Centrolenidae), in which females deposit eggs on a variety of substrates (Goyes Vallejos et al. 2024), providing a rare opportunity to examine how substrate choice influences offspring fitness.

*Espadarana prosoblepon* females do not provide parental care after egg laying, and thus, embryo survival depends entirely on the quality of the selected oviposition site (Goyes Vallejos et al. 2024). Consistent with the ‘mother knows best’ hypothesis (Refsnider and Janzen 2010), we hypothesized that *E. prosoblepon* females select oviposition sites that provide conditions favorable for embryonic development while minimizing key sources of mortality. To test this, we combined field monitoring of clutches with two-choice experiments in semi-captivity to assess the effects of substrate type on hatching success, embryonic development, and the incidence of different sources of embryo mortality, and to determine whether females actively prefer one substrate over another.

## Materials and methods

### Study site

We conducted our study at Las Cruces Biological Station (Organization for Tropical Studies), Coto Brus, Puntarenas, Costa Rica (8°47′10″N, 82°57′32″W; 1100 m a.s.l.). The rainy season spans May–November, with a mean annual precipitation of 3500–4000 mm and a mean temperature of 21°C (Zahawi et al. 2015). We established two 50 m transects in the station’s botanical garden along small watercourses draining into Culvert Creek, where the understory supports palms, aroids, gingers, and bromeliads, and riverine vegetation includes ferns, bryophytes, and spike mosses.

### Clutch surveys and substrate characterization

We conducted visual encounter surveys for *E. prosoblepon* egg clutches between June 9 and July 24, 2024, including one morning (0900 – 1100 h) and one evening (1900 – 2100 h) search.

Surveys were conducted on the watercourse for each of the 50 m transects, with thorough inspection of the surrounding vegetation on both sides of the water edge and up to 2.5 m above the water surface. Once a clutch was located, we recorded a unique identification number, developmental stage following Gosner (1960), clutch size (number of eggs), substrate type, and the number of neighboring clutches within a 20 cm radius. We also measured the distance from the watercourse edge (cm) and the height (cm) measured from either the ground or the water surface, depending on the substrate directly beneath the clutch.

Substrate types were classified into discrete categories (*sensu* Goyes Vallejos et al. 2024): the spike mosses (Selaginellaceae: Tracheophyta) (1) *Selaginella diffusa*; and (2) *S. haematodes*; (3) ‘bryophytes’ *sensu lato* (mosses and liverworts); (4) angiosperm stems; (5) fern blades (Pterophyta); (6) leaf litter; (7) mud (bare soil); (8) angiosperm roots; (9) spicules of the giant fern *Angiopteris evecta*.; and (10) angiosperm leaves (hereafter referred to as ‘leaves’). The frequency of substrate use across categories was compared with that reported in an earlier study of the same population (Goyes Vallejos et al. 2024) to identify changes in substrate-use trends over time.

### Microclimate Characterization

To assess differences in microclimatic conditions across substrates, we included only 23 egg clutches found in the field at developmental stages at or prior to Gosner stage 18 (i.e., embryos without a muscular response; Gosner 1960). Clutches at these early stages are less than ca. 6 days old, reducing the likelihood that eggs had already been predated or otherwise lost, and thus, are more likely to represent complete clutches at the time of discovery. Microclimatic variables (temperature, humidity, and rainfall) were monitored daily throughout embryonic development until all tadpoles had hatched. Because clutches were encountered at different times, monitoring periods differed among clutches. To ensure comparable environmental measurements across clutches, we restricted the data analysis to a standardized timeframe (July 1 – 12, 2024), during which all egg clutches had concurrent data. We placed a Kestrel Drop D2 datalogger next to each clutch to record temperature and humidity every 30 minutes. We positioned a 50 mL Falcon® tube adjacent to each clutch and recorded daily accumulated rainfall between 0700 and 1000 h the following morning.

Because clutches were distributed unevenly across substrate types, we restricted microclimatic comparisons to the two substrates with the largest sample sizes (*S. diffusa* and leaves), as the other substrates had insufficient sample sizes for meaningful comparisons.

Exploratory analyses indicated that mean and minimum temperatures varied little between the two substrates, whereas maximum temperature exhibited the greatest variability. For humidity, the minimum relative humidity showed the greatest variability compared to the maximum and the mean. We therefore used maximum temperature and minimum humidity to compare daily and hourly microclimatic patterns between substrates over time. We compared the clutches’ temperature and humidity time series per substrate using Euclidean distance (*D*; ‘*TSdist*’ package; Mori et al. 2016) as a dissimilarity metric. For each pair of clutches —one on *S. diffusa* and one on a leaf— *D* was calculated as the sum of the squared differences between temperature or humidity values recorded at the same time point. For daily comparisons, we used the maximum temperature or minimum relative humidity recorded each day; for hourly comparisons, we used the maximum temperature or minimum relative humidity at each 30-min interval across a standardized 24-hour cycle. Because we had multiple clutch replicates for each substrate, we calculated *D* for all possible *S. diffusa* – leaf pairwise comparisons and averaged these values to obtain a single mean observed Euclidean distance between the substrates (*D*^-^_obS_). To assess whether the observed difference was greater than expected by chance, we used a permutation test in which substrate labels were randomly reassigned across all clutch time series and calculated a random Euclidean distance (D*^-^_random_*). This process was repeated 10,000 times to generate a null distribution, and an empirical *P*-value was obtained as the proportion of *D*^-^_random_ values greater than or equal to *D*^-^_obS_ (right-tailed hypothesis test). This permutation approach accounts for the serial correlation in temperature and humidity data recorded at regular intervals.

We compared the cumulated rainfall between substrates using a Mann-Whitney U test (*‘stats’* package), calculated as the sum of daily maximum rainfall values recorded at each clutch over the same 12-day monitoring window used for temperature and humidity.

### Substrate effect on hatching success

To assess whether substrate type influences embryo fate and hatching success, we monitored the clutches *in situ*, and identified specific causes of mortality: (1) dehydration, indicated by shriveled egg jelly; (2) fungal infection, indicated by the presence of hyphae on embryos; (3) predation, indicated by missing or empty jelly capsules; (4) failure to develop, defined as embryos that ceased development relative to others in the clutch; (5) rain stripping, when clutches disappear after heavy rains (Warkentin 2000; Goyes Vallejos and Ramirez-Soto 2020); and (6) unknown, assigned when eggs disappeared and the cause could not be confidently determined. We placed a plastic container filled with creek water beneath each clutch to capture tadpoles from the onset of hatching until no further tadpoles remained in the clutch (typically eight days). All collected tadpoles were counted and released back into the stream. For each clutch, hatching success was calculated as the proportion of eggs that hatched relative to total clutch size. Similarly, for each source of embryonic mortality, we calculated the proportion of eggs lost to that cause relative to total clutch size. All proportions are reported as means ± SD unless otherwise noted, and range from 0 to 1, where 0 indicates no eggs were affected, and 1 indicates complete clutch loss.

We analyzed total hatching success for each substrate by modeling the proportion of hatched eggs as a function of substrate using a Generalized Linear Mixed Model (GLMM) with a binomial distribution, a logit link function, and clutch ID as a random effect, using the package *‘lme4’* (Bates et al. 2015). To determine whether individual sources of embryonic mortality varied between substrates, we modeled the proportion of eggs lost to each mortality cause separately as a function of substrate using beta regression in the ‘*betareg*’ package (Cribari-Neto and Zeileis 2010), which accounts for the overdispersion found in our proportion data. Because beta regression cannot handle exact zeroes or ones, proportions were rescaled to fall strictly within the range of zero and one using the standard correction proposed by Smithson and Verkuilen (2006).

Predation differed from other sources of embryonic mortality because losses tended to be all-or-none: either a clutch experienced no predation, or most of the clutch was predated. This produced a zero- and one-inflated distribution that standard GLMs handle poorly, especially given our small sample sizes. We therefore complemented our analysis with a Bayesian framework specifically for predation. We modeled the proportion of eggs predated as a function of the substrate and clutch ID as a random effect with a binomial distribution using the Stan programming language (Stan Development Team 2025) via the ‘*brms’* package (Bürkner 2017) in R. Sampling was run in four chains, each with 5000 posterior draws from priors with 10% warm-up. We obtained an effective sample size (ESS) of over 6900 for main effects. We visually diagnosed chains using the packages ‘*bayesplot*’ (Gabry and Mahr 2025) and ‘*priorsense’* (Kallioinen et al. 2023), and assessed their convergence, validating *R̂* values close to 1 (Vehtari et al. 2021). We calculated the 89% highest-density intervals (HDI) for the posterior probability estimates using the package ‘*HDInterval*’ (Meredith and Kruschke 2022). Non-overlapping HDIs between substrates indicate highly credible differences in predation probability.

### Oviposition substrate choice experiments

To test whether *E. prosoblepon* females actively choose among oviposition substrates and whether substrate type influences hatching success and embryonic development, we used an experimental approach with the two substrates most frequently used in nature: *S. diffusa* and leaves. We conducted a controlled two-choice experiment inside mesh enclosures (30 × 40 × 40 cm). The enclosures were placed within our study area and experienced the same ambient environmental conditions as natural clutches. Females were simultaneously offered a choice between *S. diffusa* and saplings of *Heliocarpus appendiculatus* (Malvaceae), both of which naturally occur along the transects in our study area. *Selaginella diffusa* was propagated clonally by inserting fragments into a soil-fertilizer mix in a 15 x 13 cm mesh wooden frame. A wide-weave metallic mesh covering the wooden frame provided support for *S. diffusa* growth. The frames were maintained under shaded, natural conditions near the creek for 14 days prior to the experiments. Saplings of *H. appendiculatus* were grown in a greenhouse until they reached approximately 25 cm in height and had at least five leaves. Each enclosure contained leaf litter, a water-filled container, and a small piece of bamboo in the corner, serving as a daytime refuge for the frogs. The two substrates were positioned on opposite sides of the enclosure, with *S. diffusa* hanging vertically to mimic its natural growth habits. In addition, we conducted a no-choice experiment using a similar setup, except that each enclosure contained only one substrate (either *S. diffusa* or *H. appendiculatus*). This design allowed us to confirm that the patterns observed in the two-choice experiments reflected female oviposition site preference.

For both experiments, we searched for amplectant pairs (males clasping females in a “mating embrace”) of *E. prosoblepon* along our transects between 1900 and 2200 h. Once a pair was located, it was carefully transported in a plastic bag to the enclosures and randomly assigned to either the two-choice or no-choice experiment. Pairs were kept overnight in the enclosures and returned to their point of capture within 24 h. Prior to release, females were photographed and marked with a Visual Implant Elastomer (Northwest Marine Technology, Inc.) to prevent recapture. For the two-choice experiment, we recorded the substrate on which each female laid her eggs; clutch size was recorded for both experiments. The following morning, oviposition substrates were transferred to a nearby greenhouse bench fitted with a vertical mesh (60 cm) along its middle. The *S. diffusa* wooden frames were suspended from the mesh at approximately one meter above the ground, while *H. appendiculatus* saplings were placed in bamboo pots on the bench. We placed water trays underneath all clutches to collect hatched tadpoles. Clutches were monitored twice daily (0700 and 1500 h) until all tadpoles hatched or died. For each clutch, we recorded hatching onset and offset (the interval between first and last tadpole hatching within a clutch), daily developmental stage (Gosner, 1960), and sources of mortality using a handheld microscope. Hatching success and individual sources of embryonic mortality in the choice experiments were analyzed using the same GLMM and beta regression approaches described above for the clutches monitored in the field.

We used a time-to-event analysis to compare embryonic developmental rates between eggs laid on *S. diffusa* and leaves from the choice experiments, with the ‘success event’ defined as reaching hatching competency (Gosner stage 25). This approach accounts for embryos that progressed through the next developmental stages and those that died before hatching. We tested for differences in developmental rates between the substrates using a log-rank test and visualized the results with Kaplan-Meier survival curves (package ‘*survminer*’; Kassambara et al. 2025). We then fitted a Cox proportional hazard model to estimate the effect of substrate on the likelihood of reaching hatching competency, expressed as a hazard ratio (HR) of *S. diffusa* relative to *H. appendiculatus*, and used the Schoenfeld test to validate model assumptions (package ‘*survival*’; Therneau 2024).

All analyses were performed in R v. 4.5.1 (R Core Team 2025) using RStudio (Posit Team 2025), with a significance threshold of *P* = 0.05. Model assumptions for the GLMMs were validated using the package *’performance’* (Lüdecke et al. 2021).

## Results

### Clutch Surveys and Substrate Characterization

In total, we found 153 *E. prosoblepon* egg clutches across our two transects. Most were laid on *S. diffusa* (n = 72), followed by bryophytes (n = 24), leaves (n = 23), and *S. haematodes* (n = 11). All other substrates were used infrequently (stem, n = 9, leaf litter, n = 6; fern blades, mud, plant roots, and *Angiopteris evecta* spicules, n = 2 each). Identified angiosperm and fern species are listed in Online Resource 1, Table S1. Substrate use was similar to that reported by Goyes Vallejos et al. (2024), with *S. diffusa* (47.1% in 2024; 30.2% in 2021) and bryophytes (15.7% in 2024; 27.9% in 2021) as the most common substrates. Leaves ranked third in 2024 (15%) and fourth in 2021 (9.3%), whereas leaf litter ranked third in 2021 (18.6%; Fig 1). Clutches were located at a mean distance of 25.6 ± 26.9 cm from the watercourse (range = 0 – 149 cm) and a mean height of 46.9 ± 9.2 cm (range = 0 – 303 cm). Of the 153 clutches, 73 (47.7%) had between one and four neighboring clutches within a 20 cm radius (median = 1), with clutches on *S. diffusa* and bryophytes showing the highest frequency of aggregation. Summaries by substrate are provided in the Online Resource 1, Table S2.

**Fig. 1.**
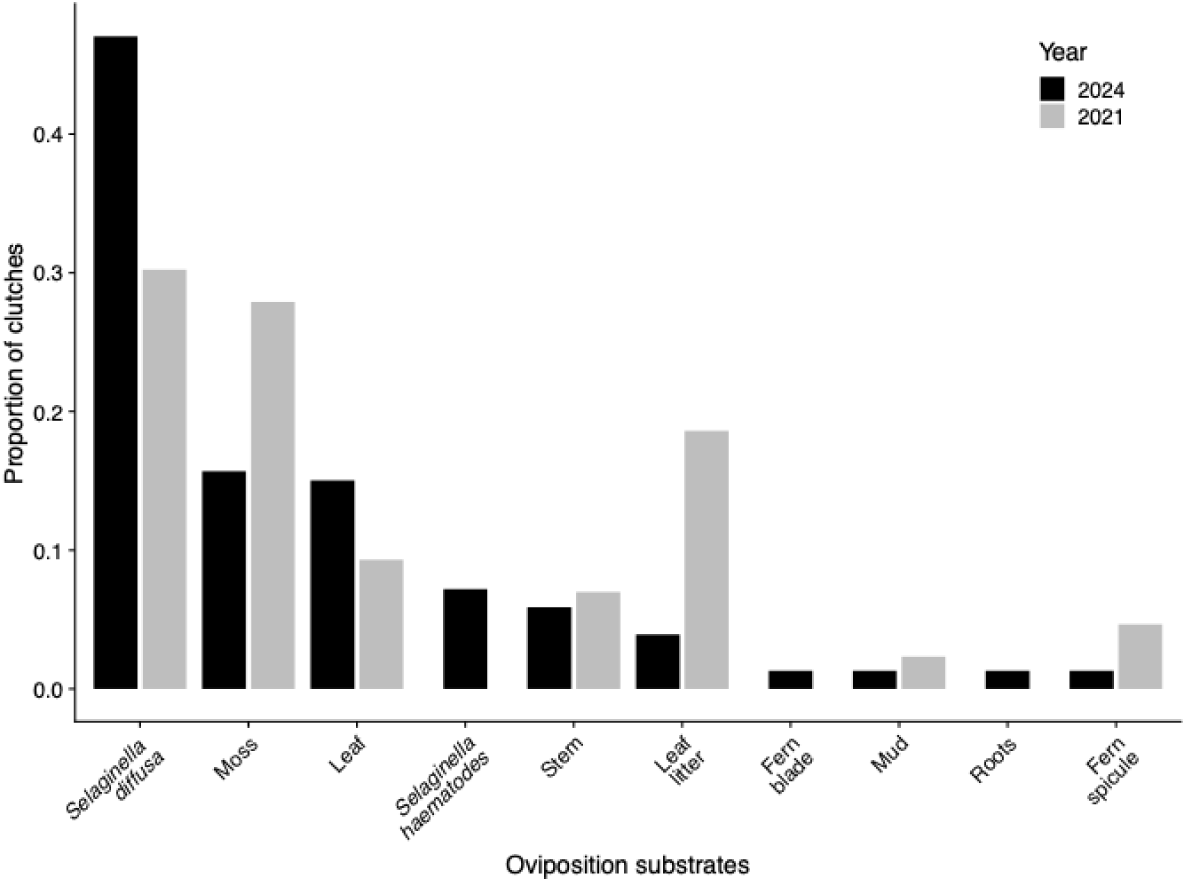
Proportion of egg clutches deposited on each substrate type recorded during two field seasons was broadly consistent between years: 2021 (Goyes Vallejos et al. 2024; n = 43) and 2024 (this study; n = 153). *Selaginella diffusa* was the most frequently used substrate in both years.

### Microclimate Characterization

The 23 monitored clutches were laid on *S. diffusa* (n = 8), leaves (n = 7), bryophytes (n = 2), *S. haematodes* (n = 3), and stems, fern blades, and plant roots (n = 1 each). The clutches on *S. diffusa* experienced mean daily temperatures of 20.6 ± 1.8 °C (range: 17.7 – 36.4), while clutche on leaves experienced mean daily temperatures of 20.3 ± 1.6 °C (range: 17.4 – 33.8, Fig. 2a). Hourly temperature values increased during the daytime, especially at noon (Fig. 2b). Neither maximum daily temperature nor hourly temperature patterns differed significantly between *S. diffusa* and leaves (daily: = 12.3, *P =* 0.54, Fig. 2e; hourly: = 11.6, *P =* 0.20, Fig. 2f). Clutches on *S. diffusa* and leaves experienced a mean daily humidity of 99.6 ± 1.5% (range: 76.4 – 100) and 99.7 ± 2.1% (range: 58.7 – 100), respectively (Fig. 2c). Across both substrates, relative humidity declined during daytime hours, particularly at noon (Fig. 2d). However, neither minimum daily humidity nor hourly humidity patterns differed significantly between *S. diffusa* and leaves (daily: = 24.0, *P =* 0.22, Fig. 2g; hourly: = 27.2, *P =* 0.57, Fig. 2h). The cumulated rainfall recorded for *S. diffusa* egg clutches (69.5.2 ± 38.9 mm, range: 4.1 – 115) did not differ significantly from that of leaves (23.0 ± 49.8.0 mm, range: 1.6 – 114; *W* = 24, *P* = 0.34). Together, this indicates that microclimatic conditions did not differ between the two substrates.

**Fig. 2.**
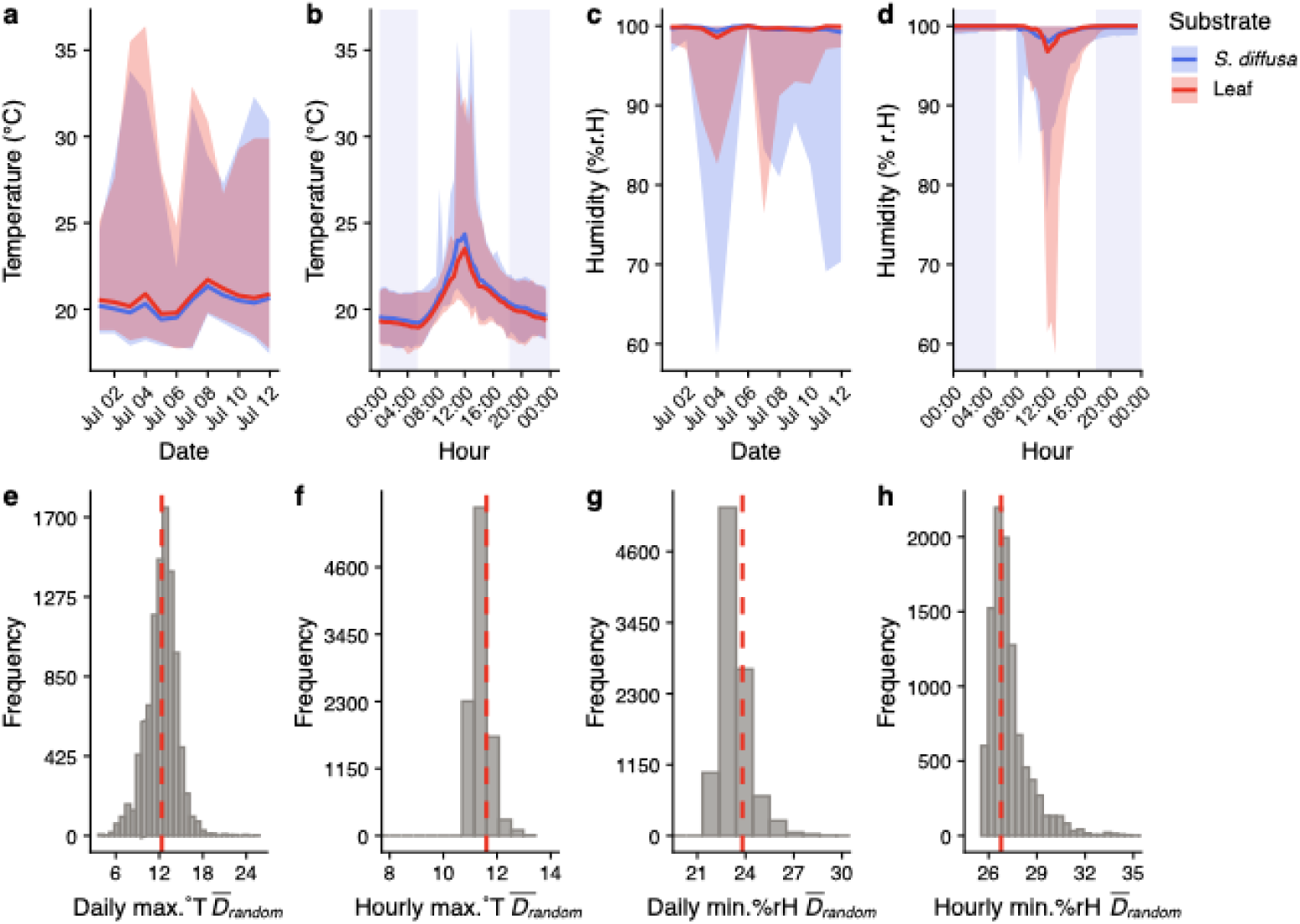
Temperature and humidity comparison between *Selaginella diffusa* and angiosperm leaves. (a) Daily and (b) hourly temperature patterns, and (c) daily and (d) hourly relative humidity patterns for each substrate, showing mean (thick line), maximum (upper line), and minimum (lower line) values; gray shaded areas in (b,d) indicate nighttime hours. Permutation tests for mean Euclidean distances in (e) daily maximum, (f) hourly maximum temperature, (g) daily minimum humidity, and (h) hourly minimum humidity. Gray bars show the null distribution (*D̄_random_*), and the red dashed line indicates the observed mean Euclidean distance (*D̄_obs_*). In all cases, *D̄_obs_* values fell within the null distribution, indicating no significant differences in maximum temperature or minimum humidity between substrates (all *P >* 0.05).

### Substrate effect on hatching success

The monitored clutches in the field had an average clutch size of 26.4 ± 4.7 eggs (range: 18 – 34, n = 23). They were found at a median Gosner stage of 13 (range: 8 – 17), began hatching at a median of 16 days after being found (range: 9 – 23), and took a median of eight additional days until hatching offset, for a median monitoring period of 25 days (range: 19 – 30). As noted, subsequent comparisons were restricted to *S. diffusa* (n = 8) and leaves (n = 7).

Hatching success was significantly lower for clutches laid on leaves (0.31 ± 0.41) than on *S. diffusa* (0.67 ± 0.23; β = -3.47, *SE* = 1.49, *Z =* -2.33, *P =* 0.02, Fig. 3a). Predation was the main source of mortality on leaves (0.53 ± 0.49) and was substantially lower on *S. diffusa* (0.048 ± 0.14, Fig. 3b). Notably, four of the seven clutches on leaves suffered complete predation (100% egg loss), two by *Polybia* sp. wasps (pers. obs) and two by unidentified predators. The Bayesian analysis confirmed that predation probability was statistically different between the two substrates, with non-overlapping 89% HDIs (Posterior probability on leaves = 0.48, HDI = 0.23 – 0.74; Posterior probability on *S. diffusa* = 0.022, HDI = 0.004 – 0.054, Fig. 4). The primary mortality source in *S. diffusa* was unknown causes (0.15 ± 0.16), which was also the second most common source on leaves (0.09 ± 0.14); this difference was not significant (β = -0.64, *SE* = 0.55, *Z =* -1.16, *P* = 0.24). We found no differences for developmental failure (0.025 ± 0.04, β = -0.095, *SE* = 0.52, *Z =* -0.18, *P* = 0.85) or rain striping (0.025 ± 0.037, β *=* 0.67, *SE* = 0.52, *Z =* 1.29, *P* = 0.19). Clutches in both substrates were free of fungal infections or embryo dehydration.

**Fig. 3.**
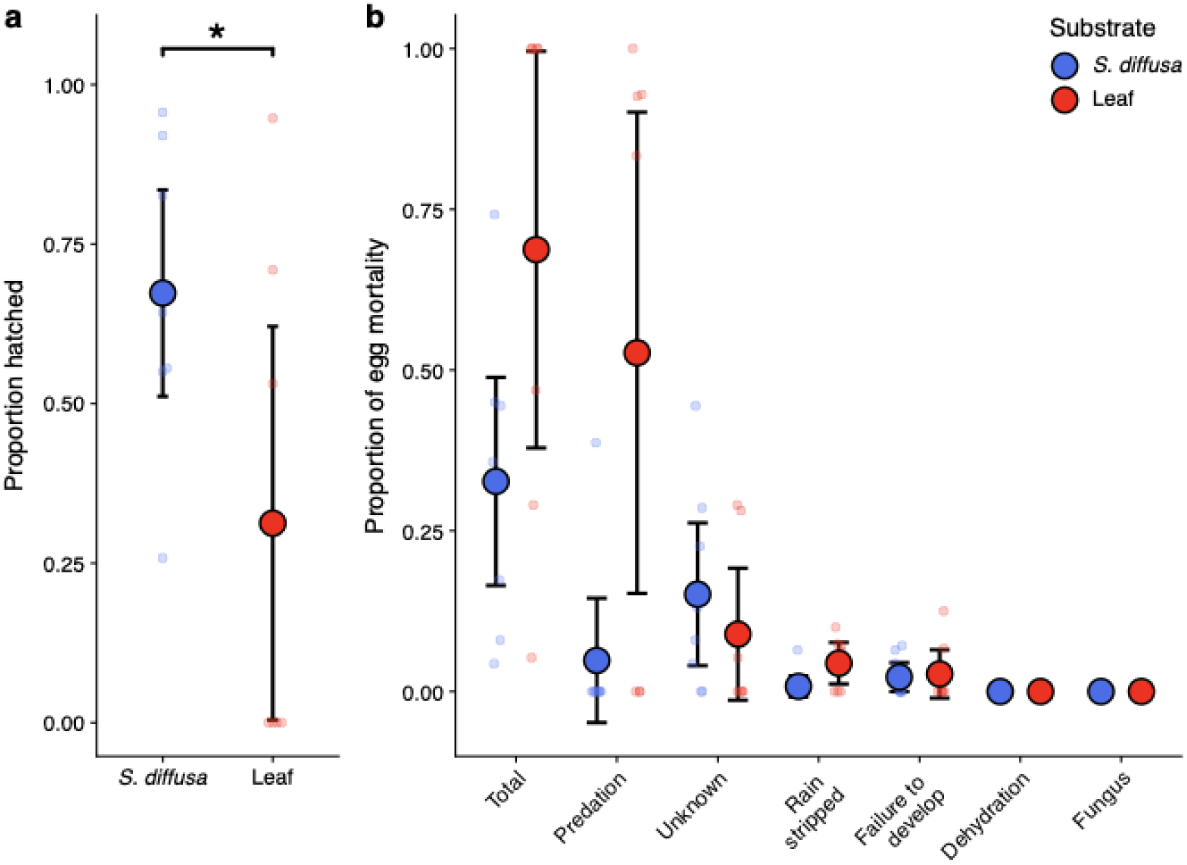
Hatching success and embryo mortality in the field. (a) Mean proportion (± SE) of eggs hatched per clutch for *S. diffusa* (n = 8) and leaf (n = 7) substrates; dots represent individual clutches. Clutches on *S. diffusa* had significantly higher hatching success than those on leaves (GLMM; * *P* = 0.02). (b) Mean proportion (± SE) of eggs lost to each source of embryonic mortality per substrate.

**Fig. 4.**
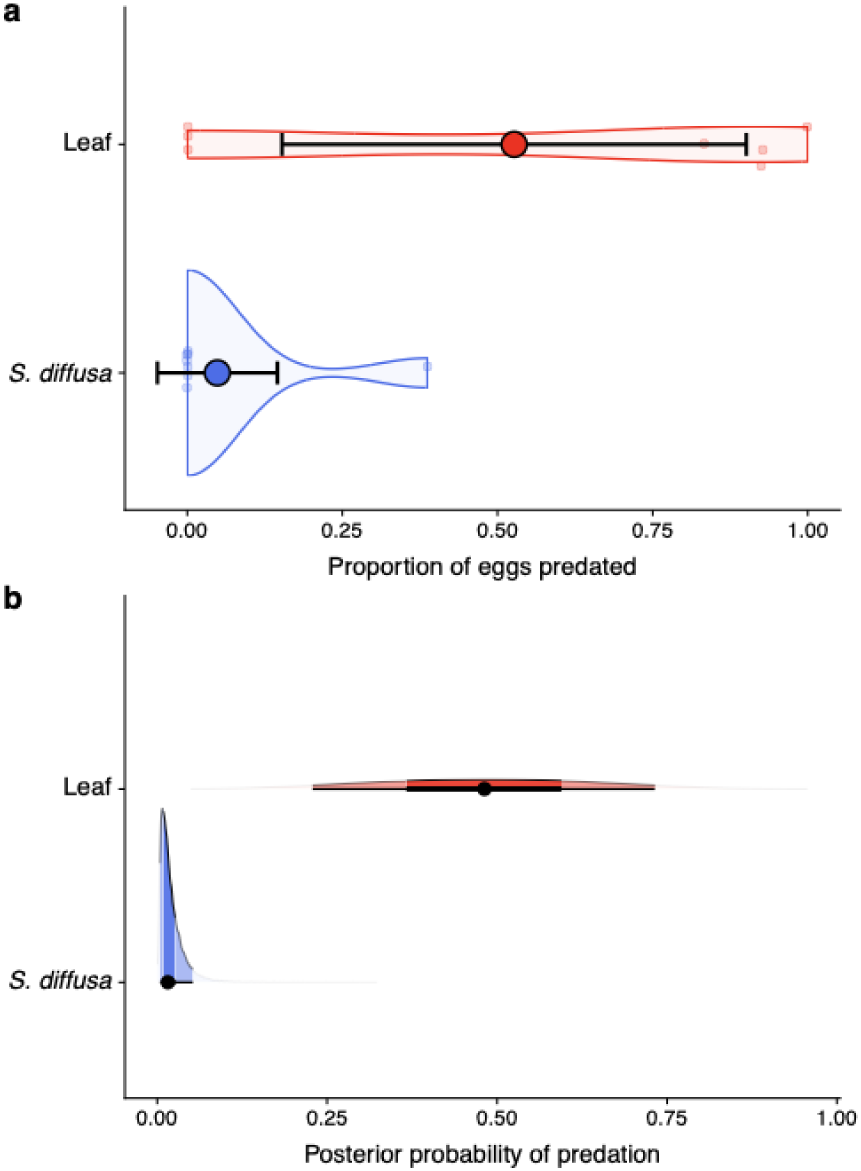
Predation probability by substrate in the field. (a) Observed proportion (±SE) of eggs predated per clutch on *S. diffusa* and angiosperm leaf substrates; dots represent individual clutches, and the shaded area shows the data distribution. (b) Posterior probability distributions of the proportion of eggs predated per substrate, estimated using a Bayesian binomial model. Dark and light bands indicate the 50% and 89% highest density intervals (HDI), respectively. Predation probability was higher on leaves (posterior mean = 0.48, 89% HDI: 0.23–0.74) than on *S. diffusa* (posterior mean = 0.022, 89% HDI: 0.004–0.054).

### Oviposition substrate choice experiments

In the two-choice experiments, all females (n = 34) laid eggs on *S. diffusa* (Fig. 5a). The average clutch size was 25.6 ± 3.87 eggs (range: 18 – 38). In the no-choice experiments, females presented with only *S. diffusa* laid their eggs on it (n = 8), whereas those presented with only leaves laid their eggs on leaves (n = 11) or on the enclosure mesh (n = 3). Clutches deposited on the mesh were excluded from the analyses. Substrate type had a significant effect on hatching success across experiments (likelihood ratio test: χ*²* = 55.35, *df* = 2, *P* < 0.0001). Hatching success for clutches on *S. diffusa* was similar between the two-choice (0.64 ± 0.25) and no-choice experiments (0.71 ± 0.23; *Z =* -1.083, *P* = 0.53), but clutches laid on leaves had markedly lower hatching success (0.024 ± 0.055), differing significantly from *S. diffusa* in both experiments (two-choice: *Z =* 7.18; no-choice: *Z =* 6.71, both *P* < 0.0001, Fig. 5b).

**Fig. 5.**
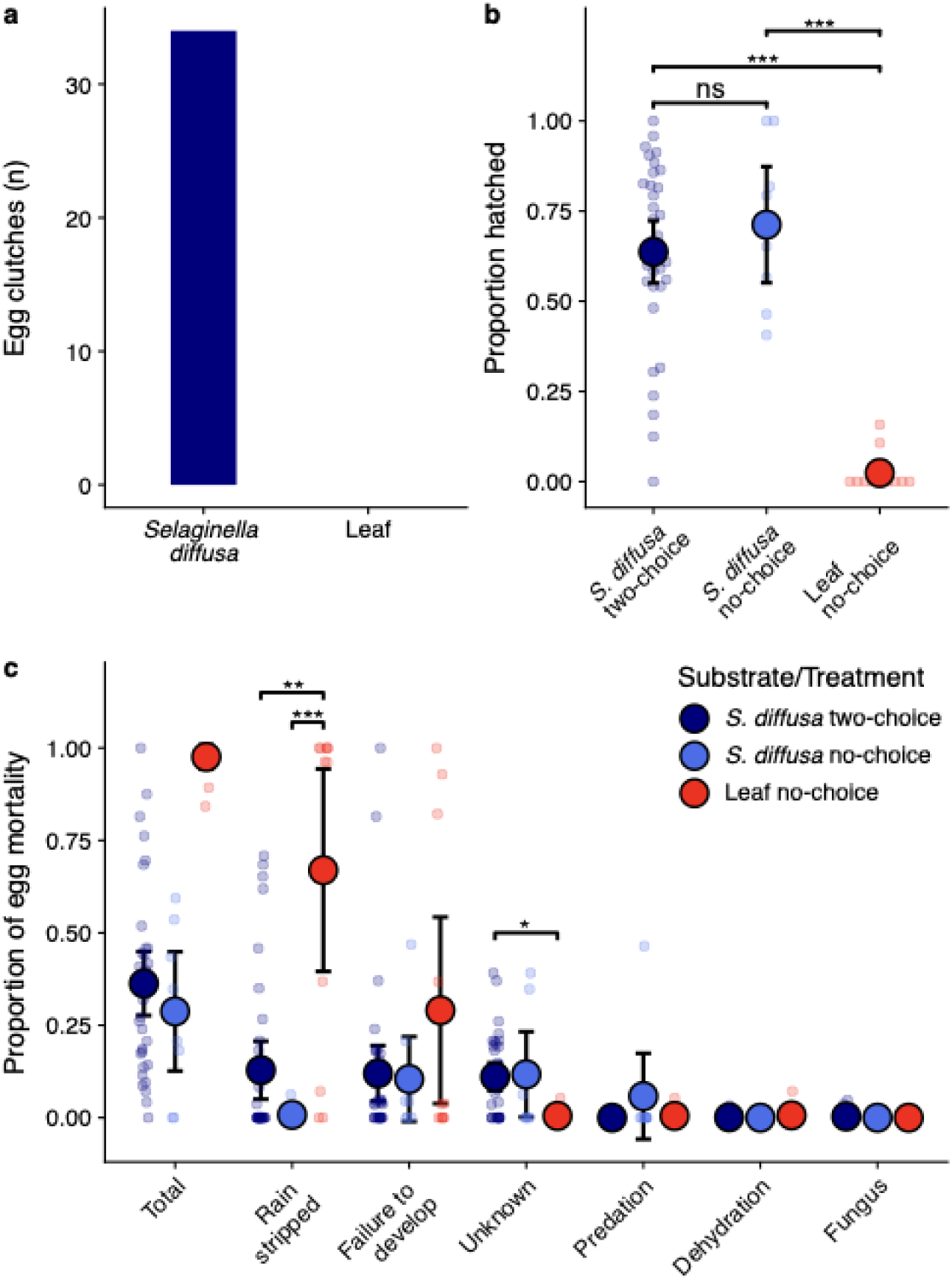
Female substrate choice, hatching success, and embryo mortality in semi-captivity. (a) All females (n = 34) deposited their clutches on S. diffusa over leaves in the two-choice experiment. (b) Mean proportion (± SE) of eggs hatched per clutch for *S. diffusa* (two-choice and no-choice) and leaf (no-choice) treatments; dots represent individual clutches. Hatching success did not differ between *S. diffusa* treatments (ns) but was significantly lower on leaves than on *S. diffusa* in both experiments (GLMM; *** *P* < 0.0001). (c) Mean proportion (± SE) of eggs lost to each source of embryonic mortality per treatment (* *P* < 0.05, ** *P* < 0.01, *** *P* < 0.0001).

Overall, embryonic mortality on *S. diffusa* was low across both experiments, with no single mortality source exceeding an average of 13% of eggs per clutch (Fig. 5c). In the two-choice experiment, the main sources of embryo mortality were rain stripping (0.13 ± 0.23), developmental failure (0.12 ± 0.22), and unknown causes (0.11 ± 0.11); no instances of predation were recorded. In the no-choice experiment, unknown causes (0.12 ± 0.16) and developmental failure (0.10 ± 0.16) were the primary sources of embryo mortality. Only one clutch suffered predation on *S. diffusa*, in which 46.4% of the eggs were lost to parasitism by *Drosophila* sp. larvae, and two embryos in another clutch were lost to rain stripping. Fungal infection and dehydration were negligible across all *S. diffusa* clutches in both experiments, with only three embryos showing signs of fungal infection and one embryo lost to dehydration in the two-choice experiment. On leaves in the no-choice experiment, rain stripping was the primary source of mortality (0.67 ± 0.45) and was significantly higher than on *S. diffusa* in both experiments (no-choice: β = -0.40, SE = 0.11, *Z = -*3.65, *P* = 0.0008; two-choice: β = -0.36, SE = 0.10, *Z = -*3.47, *P* = 0.001). Unknown causes accounted for only one embryo lost across all clutches laid on leaves, a significantly lower proportion than on *S. diffusa* in the two-choice experiment (β = 0.051, SE = 0.021, *Z* = 2.45, *P* = 0.038), but similar to *S. diffusa* in the no-choice experiment (β = 0.032, SE = 0.029, *Z* = 1.12, *P =* 0.504). All other mortality sources on leaves — developmental failure (0.29 ± 0.42), and isolated cases of dehydration (2 embryos) and predation (1 embryo)— did not differ significantly from either *S. diffusa* treatment (all *P* > 0.05). Fungal infection was not observed on leaves.

### Substrate effect on embryonic development

For the time-to-event analysis, we combined data from *S. diffusa* clutches from both experiments, as hatching success did not differ significantly between them. To minimize disturbance, embryos were counted superficially at each visit, leaving the disappearance of 96 embryos across all clutches undetected until final counts were compared with the original clutch sizes; these embryos were excluded from the analyses, as the timing of their loss could not be determined. Clutches laid on *S. diffusa* began hatching at a median of 14 – 16 days after oviposition and completed hatching at a median of 27 – 28 days (Table 1). One *S. diffusa* clutch in the two-choice experiment failed to develop entirely and was excluded from the offset calculation. On leaves, hatching onset was recorded as early as 14 days. However, hatching onset and offset could be calculated for only two of 11 clutches, as seven were entirely lost to rain stripping, and two of the remaining four experienced 100% developmental failure. In the two surviving clutches, mortality was dominated by developmental failure in one and a split between developmental failure and rain stripping in the other, each yielding only three tadpoles. These six tadpoles completed hatching at 17 and 20 days, respectively.

**Table 1.** Hatching onset and offset (days) for egg clutches across choice experiments. Values are medians with ranges in parentheses, except where noted.

| Experiment | Hatching onset (days) | Hatching offset (days) | Sample size |
| --- | --- | --- | --- |
| *S. diffusa* (two-choice) | 16 (9 – 23) | 27 (20 – 31) | 34<sup>a</sup> |
| *S. diffusa* (no-choice) | 14.5 (9 – 23) | 28 (16 – 32) | 8 |
| Leaves (no-choice) | 14, 18<sup>b</sup> | 17, 20<sup>b</sup> | 2 |

<sup>a</sup> One clutch excluded from hatching offset calculation due to complete developmental failure; n = 33 for offset.
<sup>b</sup> Onset and offset could only be determined for two clutches; individual values are reported rather than medians.

Eggs on *S. diffusa* reached hatching competency (Gosner stage 25) significantly earlier, with a median of 12 days (859 out of 985 eggs), compared to a median of 14 days for eggs laid on leaves (14 out of 278 eggs; log-rank test *P* < 0.0001). The probability of reaching hatching competency was 3.2 times higher for eggs on *S. diffusa* than on leaves (Cox proportional hazards model: *HR* = 3.12, *Z =* −4.22, *P* < 0.0001). The proportional hazards assumption was not met (Schoenfeld test: *P* = 0.007), indicating that the effect of substrate on the probability of reaching hatching competency was not constant throughout embryonic development. Developmental rates were similar between substrates early on but diverged markedly from day 11 onward (Fig. 6).

**Fig. 6.**
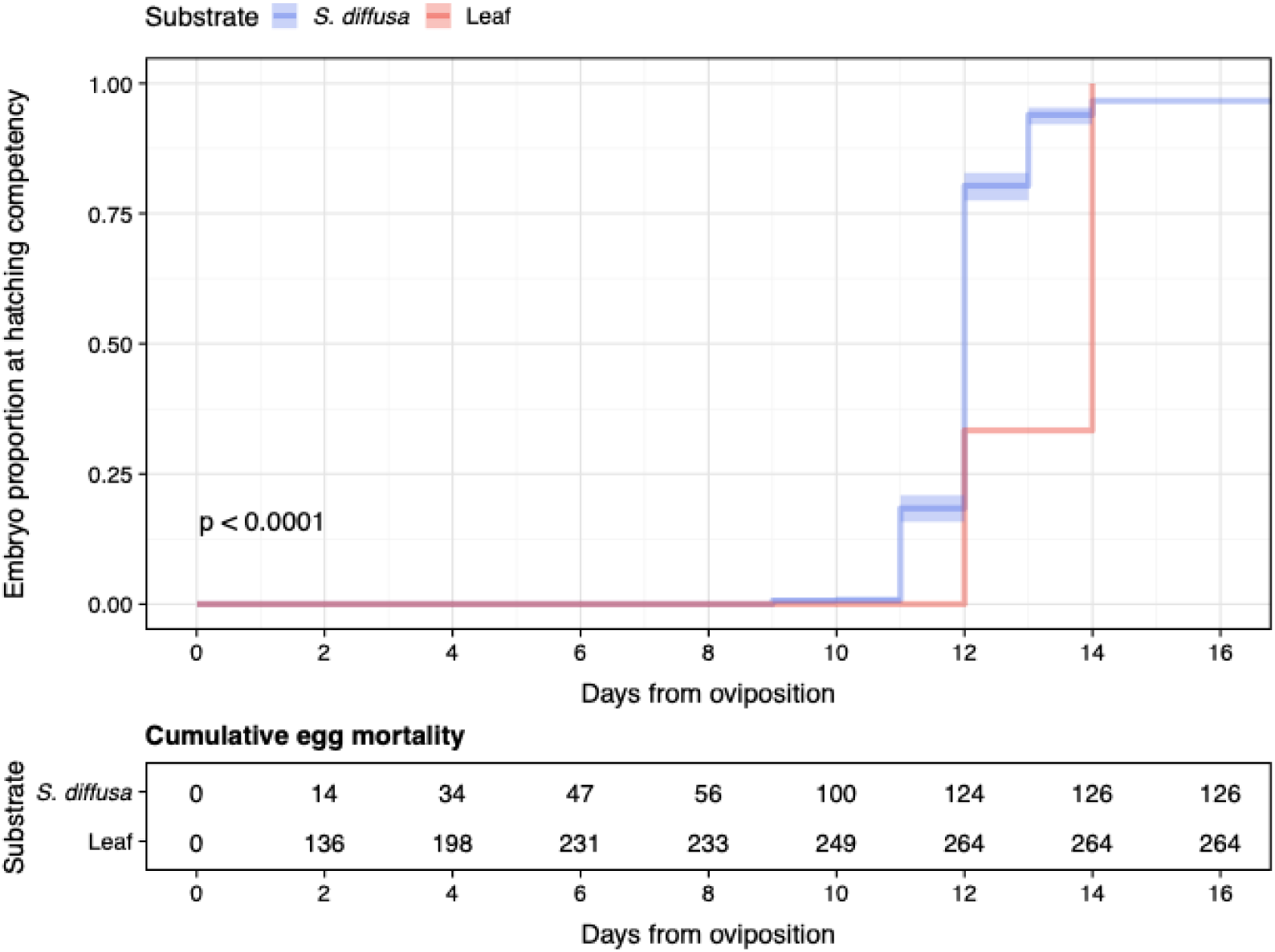
Embryonic development by substrate. Kaplan-Meier curves showing the cumulative proportion of eggs reaching Gosner stage 25 (hatching competency) over time for *S. diffusa* (n = 985 eggs) and leaf (n = 278 eggs) substrates. Shaded areas indicate 95% confidence intervals. Eggs on *S. diffusa* reached hatching competency earlier and significantly faster (median = 12 days) than those on leaves (median = 14 days; log-rank test: *P* < 0.0001). The table below shows cumulative egg mortality at each time point for each substrate.

## Discussion

Our results demonstrate that oviposition substrate has significant consequences for embryo survival in *Espadarana prosoblepon* and provide direct empirical evidence for the adaptive significance of substrate choice in an arboreal frog. Hatching success was higher on the most frequently used substrate (*S. diffusa*), where embryos experienced less predation, the primary source of embryo mortality in this species. Moreover, clutches on *S. diffusa* reached hatching competency significantly faster than those on leaves. When given a choice, females preferentially selected *S. diffusa*, linking maternal behavior directly to offspring fitness. Together, these findings support the hypothesis that non-random oviposition site selection in *E. prosoblepon* is driven by the maximization of embryo survival.

Females of *E. prosoblepon* use a variety of substrates for egg laying (Goyes Vallejos et al. 2024; this study), a behavior that contrasts sharply with the substrate specialization observed in most glass frogs, which deposit eggs almost exclusively on leaves (e.g., *H. fleischmanni*: Delia et al. 2010; *C. granulosa*: Williams 2024). Even *Espadarana andina*, the congeneric sister species, has been reported to oviposit only on the upper leaf surface (Cabanzo-Olarte et al. 2013). Despite this plasticity in oviposition substrate use, *S. diffusa* and bryophytes are consistently the most frequently used substrates across populations and years, with bryophyte use reported in two additional Costa Rican populations (Jacobson 1985; Ortiz-Ross et al. 2020). This suggests that the use of substrates such as *S. diffusa* is not merely opportunistic. The ability of females to discriminate among and select-high quality oviposition sites can have profound consequences for embryo survival. In *Drosophila melanogaster*, females shift oviposition preference toward high-ethanol fruits associated with decomposition because ingested ethanol protects embryos from parasitic wasps, even when hatching success is higher on ethanol-free substrates, revealing a fitness trade-off in oviposition site choice (Kacsoh et al. 2013). This example highlights how oviposition site selection is inherently context-dependent, shaped by local substrate availability and the specific selective pressures operating in a given environment. In *E. prosoblepon*, females lay eggs on *S. diffusa* more frequently, even where it is not the most abundant substrate (Goyes Vallejos et al. unpublished data), suggesting that they are capable of actively locating and assessing substrate quality rather than simply ovipositing on whatever is available. However, it is important to note that our study population is found in a heavily disturbed area where *S. diffusa* is present, but this substrate may be absent or rare in more pristine forest habitats across the species’ range. How females adjust their oviposition decisions in populations where *S. diffusa* is unavailable, and whether alternative substrates can provide comparable fitness benefits, is yet to be investigated. What our results do demonstrate is that females possess the capacity to discriminate among substrates and select those associated with higher offspring survival, although the sensory cues they use to locate and evaluate oviposition sites also remain to be identified. Whether females rely on visual, tactile, or chemical properties of the substrate, or on indirect cues such as the presence of conspecific clutches, represents a compelling avenue for future research, shedding light on the proximate mechanisms underlying this maternal behavior.

### Substrate effects on embryo survival and development

The differences in hatching success between substrates were unlikely to be driven by microclimatic conditions, as temperature, humidity, and rainfall did not differ between *S. diffusa* and leaves. This suggests that the characteristics of the substrate itself underlie the observed differences in embryo survival. Clutches monitored in the field had hatching success twice as high on *S. diffusa* compared to leaves, a difference driven primarily by more predation events on clutches found on leaves. We observed wasp predation directly on two clutches laid on leaves, both of which were completely decimated. Wasps extract embryos one by one and can kill an entire clutch within hours, making these predation events catastrophic once a clutch is discovered. Wasp predation on anuran eggs has been reported in other Neotropical arboreal frogs, including other centrolenids (Warkentin 2000; Touchon and Warkentin 2009; Vockenhuber et al. 2009; Chaves-Acuña et al. 2020). Seemingly, wasps actively patrol vegetation overhanging streams, where egg clutches on leaves are predictably located and relatively exposed. Once a wasp locates a clutch on a leaf, the embryos have no means of defense (Warkentin 2000).

*Selaginella diffusa* forms dense creeping mats along the walls of ditches at our study site, effectively concealing clutches from visually searching predators such as wasps. Opportunistic predators such as crickets, opilionids, and ants may occasionally encounter clutches by chance but typically cause only minor losses. The role of predation risk in shaping oviposition site selection is widespread across taxa, with natural selection favoring the ability to identify low-risk sites, detect predator cues, or adjust nest placement after offspring loss (birds: Latif et al. 2012; Mainwaring et al. 2015, reptiles: Refsnider and Janzen 2010, amphibians: Binckley and Resetarits 2002; Resetarits 2005; Touchon and Worley 2015, invertebrates: Silberbush and Blaustein 2011; Otsuki and Yano 2017). In *E. prosoblepon*, females carry the male on their back during amplexus, which begins around 1900 hours, and move several meters throughout the night before settling on an oviposition site, with most egg laying occurring between midnight and 0300 hours (Goyes Vallejos et al. 2024). Whether these nocturnal excursions reflect actively searching for a preferred substrate or serve other functions remains unknown, but the strong preference for *S. diffusa* demonstrated in our two-choice experiment suggests that oviposition site choice in this species is far from indiscriminate.

Beyond differences in hatching success, clutches on *S. diffusa* also showed faster embryonic development. Embryos on this substrate reached hatching competency (Gosner stage 25) earlier than those on leaves. Gosner stage 25 marks the point at which tadpoles are capable of free swimming, active feeding, and survival outside the egg capsule (Gosner 1960), and thus represents a critical developmental threshold. Interestingly, reaching hatching competency does not necessarily lead to immediate hatching. Experimental clutches on both substrates completed hatching between 20 and 30 days; thus, embryos remain in the egg for several days after becoming capable of hatching. A potential explanation for this delay is that embryos benefit from a prolonged growth period. Prematurely hatched *E. prosoblepon* embryos exhibited clumsy movements and reduced activity compared to those that had reached full competency (pers. obs.). Delaying hatching beyond this threshold may therefore allow for further locomotor and digestive system development, improving swimming performance and foraging efficiency after entering the aquatic environment (Delia et al. 2019), as well as enabling tadpoles to synchronize hatching with favorable environmental conditions, such as heavy rainfall that flushes tadpoles into the water. Similar strategies have been documented in other taxa, such as the California grunion (*Leuresthes tenuis*), whose embryos delay hatching until the tide reaches the eggs buried in the sand, reducing desiccation risk (Smyder and Martin 2002). Consequently, by reaching hatching competency earlier on *S. diffusa*, embryos gain additional days to maximize development before hatching, potentially improving their survival and performance in the aquatic habitat.

One notable finding in the choice experiments was the high incidence of rain stripping on leaf clutches, which contrasts with field observations from this study and Goyes Vallejos et al. (2024), where rain stripping was a minor source of mortality. This discrepancy likely reflects an, experimental artifact, similar to what was observed in a study by Goyes Vallejos and Ramirez-Soto (2020), where rain stripping emerged as a significant mortality source in an experiment in which pairs in amplexus were allowed to oviposit on fern fronds inside cages, which were then manually placed in the forest. Both the manipulation and the substrate type used in these experimental approaches may have contributed to detachment. Females of *E. prosoblepon* use a diversity of plant species as oviposition substrates in the field; however, no single species — apart from *Crinum americanum*, which was locally abundant in one of our transects— was used more than a couple of times, and it remains unknown whether *H. appendiculatus*, the species used in our experiment, is naturally selected by females in the field. Because our experiment relied on a single leaf species, we cannot determine whether the high rain stripping observed reflects a general property of leaves as a substrate category or is specific to the physical characteristics of *H. appendiculatus*, and thus, whether differences in rain stripping susceptibility exist among the many plant species used naturally by *E. prosoblepon* females. The leaves used in the experiment may have been particularly susceptible to rain stripping, since we used saplings with flexible leaves, and clutches can become engorged and heavy enough to detach from the leaf surface after the first rains since oviposition. This can occur on *S. diffusa* as well, though to a lesser extent, as clutches tend to become entangled in the stems and leaves, providing additional mechanical support. This experimental artifact may have exaggerated the apparent disadvantage of leaves as an oviposition substrate, and future experiments should carefully consider leaf species and physical characteristics to better reflect natural oviposition choices.

Despite the clear fitness advantages of *S. diffusa*, females continue to oviposit on leaves and other substrates in the wild. Due to sample size limitations, we could not fully assess hatching success across all substrates used by *E. prosoblepon* females, but the persistence of leaf use raises an important question: why does oviposition on leaves persist in this population despite its apparent costs? Habitat variability may create conditions in which the relative quality of substrates fluctuates across seasons or years, thereby maintaining the use of multiple substrates as an adaptive strategy (Devries et al. 2018). Indeed, the capacity to use a broad range of oviposition sites may itself be the trait under selection, providing a buffer against the loss or reduced availability of any single preferred substrate. For instance, in the cotton bollworm *Helicoverpa armigera*, females can oviposit on dozens of plant species and readily switch to alternative substrates when preferred host plants are unavailable (Nataraj et al. 2024).

Alternatively, if the heritability of oviposition site preference is low or absent, as has been suggested for nest-site choice in painted turtles (*Chrysemys picta;* Topping and Valenzuela 2021), then selection cannot efficiently fix a preference for *S. diffusa* in the population, and leaf use may persist simply because the behavioral tendency to choose *S. diffusa* cannot be reliably passed from mother to offspring. In *E. prosoblepon*, a broad substrate repertoire may therefore reflect both habitat variability and a generalist reproductive ecology in which plasticity, rather than strict preference, is the adaptive solution. Distinguishing among these possibilities will require long-term data on substrate availability, female preference, and fitness outcomes across populations and seasons.

## Conclusions

Our findings have broad implications for understanding oviposition site selection in arboreal frogs. Most studies investigating oviposition site selection rarely distinguish between substrate use driven by availability and active preference driven by fitness consequences. Here, we provide empirical evidence that females of *E. prosoblepon* not only use *S. diffusa* more frequently than other substrates, but actively prefer it when given a choice, and that this preference translates into measurable fitness benefits through reduced predation and higher hatching success, consistent with the hypothesis that maternal oviposition site selection is driven by the maximization of embryo survival (Refsnider and Janzen 2010). Future studies should examine how substrate use and preference vary across seasons, as the dry and wet seasons likely differ in predator communities, rainfall intensity, or substrate availability, potentially shifting the relative quality of oviposition substrates. Comparisons across populations in different habitats would further clarify whether the preference for *S. diffusa* or functionally similar substrates is a fixed trait or a context-dependent response to local conditions. Ultimately, our study highlights the importance of integrating behavioral, experimental, and field approaches to understand the adaptive significance of maternal behaviors related to habitat choice for offspring survival.

## Supporting information

Online Resource

## Acknowledgments

We thank the Organization for Tropical Studies and the staff at Las Cruces Biological Station for logistical support, especially to Rodolfo Quiros. We thank Michael Atencio for preparing the frames, cultivating *S. diffusa* for our experimental setup, and providing additional logistical support. We are grateful to Lilly Briggs and David Rodríguez (Finca Cántaros Environmental Association) for providing *H. appendiculatus* plants for the two-choice experiments, and to Jeffrey Flores for identifying the plant species used for oviposition. Natalia Mejía and Alma Jarbou assisted during field surveys. We also thank Kevin Middleton, Rex Cocroft, and Manuel Leal for their valuable feedback on the experimental design, analyses, and manuscript.

## Ethics approval

All behavioral observations and field manipulations followed the Association for the Study of Animal Behaviour and the Animal Behavior Society Guidelines for the treatment of animals in behavioral research and teaching. Our study was approved by the Costa Rican Ministry of the Environment and Energy (MINAE) and the National System of Conservation Areas (SINAC) (approval numbers: R-SINAC-PNI-ACLAP-029-2024 and SINAC-ACLAP-DR-GASP-PNI-R-0019-2025), as well as the Animal Care and Use Committee at the University of Missouri (Protocol No. 45158).

## Author contributions

The study was designed and conceived by JGV, with contributions from JSCF. JSCF performed data collection, formal analysis, visualization, figure preparation, and data curation. The first draft of the manuscript was written by JSCF under the supervision of JGV, who subsequently reviewed and edited earlier versions. All authors read and approved the final manuscript. JGV contributed to resources, supervision, project administration, and funding acquisition.

## References

Amat JA, Masero JA (2004) Predation risk on incubating adults constrains the choice of thermally favourable nest sites in a plover. Anim Behav 67:293–300 doi: 10.1016/j.anbehav.2003.06.014

Bates D, Maechler M, Bolker B, Walker S (2015) Fitting linear mixed-effects models using lme4. J Stat Softw 67:1–48 doi: 10.18637/jss.v067.i01

Binckley CA, Resetarits WJ (2002) Reproductive decisions under threat of predation: squirrel treefrog (*Hyla squirella*) responses to banded sunfish (*Enneacanthus obesus*). Oecologia 130:157–161 doi: 10.1007/s004420100781

Bürkner PC (2017) brms: An R package for Bayesian multilevel models using Stan. J Stat Softw 80:1–28 doi: 10.18637/jss.v080.i01

Buxton VL, Sperry JH (2017) Reproductive decisions in anurans: a review of how predation and competition affects the deposition of eggs and tadpoles. BioScience 67:26–38 doi: 10.1093/biosci/biw149

Cabanzo-Olarte LC, Ramírez-Pinilla MP, Serrano-Cardozo VH (2013) Oviposition, site preference, and evaluation of male clutch attendance in *Espadarana andina* (Anura: Centrolenidae). J Herpetol 47:314–320 doi: 10.1670/11-266

Chaves-Acuña W, Salazar-Zúñiga JA, Chaves G (2020) Egg clutch survival under prolonged paternal care in a glass frog, Hyalinobatrachium talamancae. Ichthyol Herpetol 108:514–521 doi: 10.1643/CH-19-331

Cribari-Neto F, Zeileis A (2010) Beta regression in R. J Stat Softw 34:1–24 doi:10.18637/jss.v034.i02

Delia J, Cisneros-Heredia DF, Whitney J, Murrieta-Galindo R (2010) Observations on the reproductive behavior of a Neotropical glassfrog, Hyalinobatrachium fleischmanni (Anura: Centrolenidae). S Am J Herpetol 5:1–12 doi: 10.2994/057.005.0101

Delia J, Rivera-Ordonez JM, Salazar-Nicholls MJ, Warkentin KM (2019) Hatching plasticity and the adaptive benefits of extended embryonic development in glassfrogs. Evol Ecol 33:37–53 doi: 10.1007/s10682-018-9963-2

Devries, JH, Clark RG, Armstrong LM. (2018) Dynamics of habitat selection in birds: adaptive response to nest predation depends on multiple factors. Oecologia 187:305–318. 10.1007/s00442-018-4134-2

Gabry J, Mahr T (2025) bayesplot: Plotting for Bayesian models. R package version 1.14.0. https://mc-stan.org/bayesplot/

Gosner KL (1960) A simplified table for staging anuran embryos and larvae with notes on identification. Herpetologica 16:183–190

Goyes Vallejos J, Ramirez-Soto K (2020) Causes of embryonic mortality in *Espadarana prosoblepon* (Anura: Centrolenidae) from Costa Rica. Phyllomedusa 19:83–92 doi: 10.11606/issn.2316-9079.v19i1p83-92

Goyes Vallejos J, Siles JS, Calero V, Rodriguez N, Machado G (2024) Not enough time: short-term female presence after oviposition does not improve egg survival in the emerald glass frog. Anim Behav 213:161–171 doi: 10.1016/j.anbehav.2024.05.008

Gripenberg S, Mayhew PJ, Parnell M, Roslin T (2010) A meta-analysis of preference-performance relationships in phytophagous insects. Ecol Lett 13:383–393 doi: 10.1111/j.1461-0248.2009.01433.x

Haddad CFB, Sawaya RJ (2000) Reproductive modes of Atlantic Forest hylid frogs: a general overview and the description of a new mode. Biotropica 32:862–871 doi: 10.1111/j.1744-7429.2000.tb00624.x

Ichioka Y, Kajimura H (2024) Arboreal or terrestrial: oviposition site of *Zhangixalus* frogs affects the thermal function of foam nests. Ecol Evol 14:e10926 doi: 10.1002/ece3.10926

Jacobson SK (1985) Reproductive behavior and male mating success in two species of glass frogs (Centrolenidae). Herpetologica 41:396–404

Jones LC (2022) Insects allocate eggs adaptively according to plant age, stress, disease or damage. Proc R Soc B 289:20220831 doi: 10.1098/rspb.2022.0831

Kacsoh BZ, Lynch ZR, Mortimer NT, Schlenke TA (2013) Fruit flies medicate offspring after seeing parasites. Science 339:947–950 doi: 10.1126/science.1229625

Kallioinen N, Paananen T, Bürkner PC, Vehtari A (2023) Detecting and diagnosing prior and likelihood sensitivity with power-scaling. Stat Comput 34:57 doi: 10.1007/s11222-023-10366-5

Kassambara A, Kosinski M, Biecek P (2025) survminer: Drawing survival curves using ’ggplot2’. R package version 0.5.1. https://CRAN.R-project.org/package=survminer

Latif QS, Heath SK, Rotenberry JT (2012) How avian nest site selection responds to predation risk: testing an ‘adaptive peak hypothesis’. J Anim Ecol 81:127–138 doi: 10.1111/j.1365-2656.2011.01895.x

Liao WB, Lu X (2010) Breeding behaviour of the Omei tree frog *Rhacophorus omeimontis* (Anura: Rachophoridae) in a subtropical montane region. J Nat Hist 44:2929–2940 doi: 10.1080/00222933.2010.502594

Lüdecke D, Ben-Shachar MS, Patil I, Waggoner P, Makowski D (2021) performance: An R package for assessment, comparison and testing of statistical models. J Open Source Softw 6(60):3139 doi: 10.21105/joss.03139

Mainwaring MC, Reynolds SJ, Weidinger K (2015) The influence of predation on the location and design of nests. In: Deeming DC, Reynolds SJ (eds) Nests, eggs, and incubation: new ideas about avian reproduction. Oxford University Press, pp 50–64 10.1093/acprof:oso/9780198718666.003.0005

Meredith M, Kruschke J (2022) HDInterval: Highest (Posterior) Density Intervals. R package version 0.2.4. https://CRAN.R-project.org/package=HDInterval

Mori U, Mendiburu A, Lozano JA (2016) Distance measures for time series in R: The TSdist package. R package version 3.7. https://CRAN.R-project.org/package=TSdist

Nataraj N, Hansson BS, Knaden M (2024) Learning-based oviposition constancy in insects. Front Ecol Evol 12:1351400 doi: 10.3389/fevo.2024.1351400

Ortiz-Ross X, Thompson ME, Salicetti-Nelson E, Vargas-Ramírez O, Donnelly MA (2020) Oviposition site selection in three glass frog species. Ichthyol Herpetol 108:333–340 doi: 10.1643/CE-19-243

Otsuki H, Yano S (2017) Within-patch oviposition site shifts by spider mites in response to prior predation risks decrease predator patch exploitation. Ethology 123:453–459

Poo S, Bickford DP (2013) The adaptive significance of egg attendance in a Southeast Asian tree frog. Ethology 119:671–679 doi: 10.1111/eth.12108

Posit team (2025) RStudio: Integrated Development Environment for R. Posit Software, PBC, Boston, MA. http://www.posit.co/

Pruett JE, Fargevieille A, Warner DA (2020) Temporal variation in maternal nest choice and its consequences for lizard embryos. Behav Ecol 31:902–910 doi: 10.1093/beheco/araa032

R Core Team (2025) R: A language and environment for statistical computing. R Foundation for Statistical Computing, Vienna. https://www.R-project.org/

Rahman MM, Zalucki MP, Furlong MJ (2019) Diamondback moth egg susceptibility to rainfall: effects of host plant and oviposition behavior. Entomol Exp Appl 167:701–712 doi: 10.1111/eea.13338

Refsnider JM, Janzen FJ (2010) Putting eggs in one basket: ecological and evolutionary hypotheses for variation in oviposition-site choice. Annu Rev Ecol Evol Syst 41:39–57 doi: 10.1146/annurev-ecolsys-102209-144712

Resetarits Jr WJ (2005) Habitat selection behaviour links local and regional scales in aquatic systems. Ecol Lett 8:480–486 doi: 10.1111/j.1461-0248.2005.00747.x

Sánchez-Ochoa DJ, Pérez-Mendoza HA, Charruau P (2020) Oviposition site selection and conservation insights of two tree frogs (*Agalychnis moreletii* and *A. callidryas*). S Am J Herpetol 17:17–28 doi: 10.2994/SAJH-D-17-00103.1

Silberbush A, Blaustein L (2011) Mosquito females quantify risk of predation to their progeny when selecting an oviposition site. Funct Ecol 25:1091–1095 doi: 10.1111/j.1365-2435.2011.01873.x

Smithson M, Verkuilen J (2006) A better lemon squeezer? Maximum-likelihood regression with beta-distributed dependent variables. Psychol Methods 11:54 doi: 10.1037/1082-989X.11.1.54

Smyder EA, Martin KLM (2002) Temperature effects on eggs survival and hatching during the extended incubation period of California grunion, *Leuresthes tenuis*. Copeia 2002:313–320 doi: 10.1643/0045-8511(2002)002[0313:TEOESA]2.0.CO;2

Stan Development Team (2025) RStan: the R interface to Stan. R package version 2.32.7. https://mc-stan.org/

Therneau T (2024) A package for survival analysis in R. R package version 3.8–3. https://CRAN.R-project.org/package=survival

Topping NE, Valenzuela N (2021) Turtle nest-site choice, anthropogenic challenges, and evolutionary potential for adaptation. Front. Ecol. Evol. 9:808621. doi: 10.3389/fevo.2021.808621

Touchon JC, Warkentin KM (2010) Short- and long-term effects of the abiotic egg environment on viability, development and vulnerability to predators of a Neotropical anuran. Funct Ecol 24:566–575 doi: 10.1111/j.1365-2435.2009.01650.x

Touchon JC, Worley JL (2015) Oviposition site choice under conflicting risks demonstrates that aquatic predators drive terrestrial egg-laying. Proc R Soc B 282:20150376 doi: 10.1098/rspb.2015.0376

Vehtari A, Gelman A, Simpson D, Carpenter B, Bürkner PC (2021) Rank-normalization, folding, and localization: An improved R-hat for assessing convergence of MCMC (with discussion). Bayesian Anal 16:667–718 doi: 10.1214/20-BA1221

Vockenhuber EA, Hödl W, Amézquita A (2009) Glassy fathers do matter: egg attendance enhances embryonic survivorship in the glass frog *Hyalinobatrachium valerioi*. J Herpetol 43:340–344

Vonesh JR (2000) Dipteran predation on the arboreal eggs of four *Hyperolius* frog species in western Uganda. Copeia 2000:560–566 doi: 10.1643/0045-8511(2000)000[0560:DPOTAO]2.0.CO;2

Warkentin KM (2000) Wasp predation and wasp-induced hatching of red-eyed treefrog eggs. Anim Behav 60:503–510 doi: 10.1006/anbe.2000.1508

Warner DA, Andrews RM (2002) Nest-site selection in relation to temperature and moisture by the lizard *Sceloporus undulatus*. Herpetologica 58:399–407

Williams S (2024) Habitat and oviposition site preference of glass frogs (Anura: Centrolenidae) on Río Antón in El Valle de Antón, Coclé, Panamá. Panama: Tropical Ecology, Marine Ecosystems, and Biodiversity Conservation 1.

Zahawi RA, Duran G, Kormann U (2015) Sixty-seven years of land-use change in southern Costa Rica. PLOS One 10:e0143554 doi: 10.1371/journal.pone.014355

