## Supplementary material for "Female oviposition site selection influences hatching success and embryonic development in a Neotropical glass frog": Online Resource

**Electronic Supplementary Material**

**Supplementary Table 1. Angiosperm and fern species used as oviposition substrates by *Espadarana prosoblepon*.** Clutches found between June and July, 2024 at Las Cruces Biological Station, categorized by substrate type (leaves, ferns, and stems).

| Substrate | Family | Species | Count |
|---|---|---|---|
| Leaves | Amaryllidaceae | *Crinum americanum* | 11 |
| | Araceae | *Stenospermation* sp. | 3 |
| | Araceae | *Syngonium* sp. | 1 |
| | Araceae | (unidentified species) | 1 |
| | Dipentodontaceae | *Perrottetia longistylis* | 2 |
| | Cyclanthaceae | *Cyclanthus bipartitus* | 1 |
| | Melastomataceae | *Conostegia* sp. | 1 |
| | Piperaceae | *Manekia naranjoana* | 1 |
| | (Unidentified sapling) | — | 2 |
| Ferns | Athyriaceae | *Diplazium* sp. | 2 |
| Stems | Oxalidaceae | *Biophytum dendroides* | 8 |
| | Acanthaceae | *Fittonia* sp. | 1 |

**Supplementary Table 2.** Summary of clutch characteristics by substrate (mean ± SD, range). Percentages reflect the proportion of clutches within each substrate that had one or more neighboring clutches within 20 cm.

| Substrate | Number of clutches | Distance to stream edge (cm) | Height (cm) | (%) Neighboring clutches)* | | |
|---|---|---|---|---|---|---|
| | | | | 0 | 1 | 2 – 4 |
| *Selaginella diffusa* | 72 | 19.1 ± 15 (0 – 69) | 34.2 ± 35.6 (0 – 118) | 41.7 (30) | 33.3 (24) | 25 (18) |
| Bryophytes | 24 | 41.8 ± 45 (0 – 148) | 74.8 ± 63.6 (12 – 228) | 45.8 (11) | 33.3 (8) | 20.8 (5) |
| Leaves | 23 | 26.2 ± 24.9 (0 – 80) | 56.6 ± 63 (1 – 303) | 82.6 (19) | 17.4 (4) | — |
| *Selaginella haematodes* | 11 | 27.2 ± 22.8 (3 – 76) | 47.7 ± 30.3 (16 – 96) | 54.5 (6) | 18.2 (2) | 27.3 (3) |
| Stems | 9 | 40.3 ± 42.5 (8 – 149) | 50.1 ± 46.1 (0 – 112) | 22.2 (2) | 66.7 (6) | 11.1 (1) |
| Leaf litter | 6 | 17 ± 8.9 (4 – 30) | 13 ± 14.1 (1 – 40) | 83.3 (5) | 16.7 (1) | — |
| Fern blade | 2 | 39, 43 | 18, 73 | 100 (2) | — | — |
| Mud | 2 | 20, 36 | 0, 4 | 100 (2) | — | — |
| Roots | 2 | 0, 22 | 72, 99 | 50 (1) | 50 (1) | — |
| *Angiopteris evecta* spicule | 2 | 0, 23 | 130, 168 | 100 (2) | — | — |
| **Total** | **153** | **25.6 ± 26.9 (0 – 149)** | **46.9 ± 49.2 (0-303)** | **52.3 (80)** | **30.1 (46)** | **17.6 (27)** |

* Parenthetical values indicate the number of clutches.
